# The Impact of a Wireless Audio System on Communication in Robotic-Assisted Laparoscopic Surgery: A Prospective Controlled Trial

**DOI:** 10.1101/701078

**Authors:** Ziv Tsafrir, Kirsten Janosek-Albright, Joelle Aoun, Mireya Diaz-Insua, Abd-El-Rahman Abd-El-Barr, Lauren Schiff, Shobhana Talukdar, Mani Menon, Adnan Munkarah, Evan Theoharis, David Eisenstein

**Affiliations:** Minimally Invasive Gynecologic Surgery, Women’s Health Services, Henry Ford Hospital, Detroit, Michigan; Vattikuti Urology Institute, Henry Ford Hospital, Detroit, Michigan; Department of Obstetrics and Gynecology, Kaplan medical Center, Rehovot, Affiliated to the Faculty of Medicine, the Hebrew University, Jerusalem, Israel

**Keywords:** Wireless headset, communication, teamwork, robotic surgery, noise

## Abstract

**Background:** Robotic surgery presents a challenge to effective teamwork and communication in the operating theatre (OR). Our objective was to evaluate the effect of using a wireless audio headset device on communication, efficiency and patient outcome in robotic surgery.

**Methods and findings:** A prospective controlled trial of team members participating in gynecologic and urologic robotic procedures between January and March 2015. In the first phase, all surgeries were performed without headsets (control), followed by the intervention phase where all team members used the wireless headsets. Noise levels were measured during both phases. After each case, all team members evaluated the quality of communication, performance, teamwork and mental load using a validated 14-point questionnaire graded on a 1-10 scale. Higher overall scores indicated better communication and efficiency. Clinical and surgical data of all patients in the study were retrieved, analyzed and correlated with the survey results.

The study included 137 procedures, yielding 843 questionnaires with an overall response rate of 89% (843/943). Self-reported communication quality was better in cases where headsets were used (113.0 ± 1.6 vs. 101.4 ± 1.6; p < .001). Use of headsets reduced the percentage of time with a noise level above 70 dB at the console (8.2% ± 0.6 vs. 5.3% ± 0.6, p < .001), but had no significant effect on length of surgery nor postoperative complications.

**Conclusions:** The use of wireless headset devices improved quality of communication between team members and reduced the peak noise level in the robotic OR.

## Introduction

There is strong evidence in the literature that supports the importance of effective communication and teamwork in regards to patient safety in the operating room (OR) (1, 2). Event analysis has found that deficiencies in teamwork and communication contribute to adverse events, thus demonstrating that nontechnical skills are as important as technical surgical skills in preventing adverse patient outcomes (3, 4).

Communication is defined as “a process by which information is exchanged between individuals through a common system of symbols, signs, or behavior” (5). Communication in the OR relies heavily on speech, but also encompasses visual and physical cues. Due to the large footprint platform of the robot, team members are physically separated in space, and thus face-to-face interaction during the surgery is severely limited. Unlike the conventional OR setting, robotic surgeons must rely primarily on auditory means of communication, unaided by visual cues (6-8). In addition, studies have shown that increased noise during surgery was associated with a greater risk for postoperative complications (9, 10). The robotic platform possibly lead to increased noise in the OR, which could present a source for errors and impaired safety and efficiency during surgery. As the adoption of robotic surgery is expanding at a rapid rate, these concerns become relevant and may present an unforeseen source of complications.

Our team sought a novel technological solution that has been used in other settings to improve auditory communication. The primary objective of our study was to determine whether a wireless audio headset improves the quality of communication in robotic surgery. Secondary objectives were to assess the impact of using such device on the noise level during robotic procedures, efficiency and patient outcomes.

## Materials and Methods

This study design was a non-randomized, prospective controlled trial. The study was approved by the institutional review board. Patients included in the study had surgery performed by the Departments of Gynecology and Urology in the Henry Ford Health System, a large vertically integrated health system in southeast Michigan. The most common robotic procedures performed by each department were included in the study, which were the following: total hysterectomy, prostatectomy and nephrectomy.

Study participants belonged to the following OR team categories: surgeons, surgeons’ assistants (including fellows, residents and certified first assistants), surgical technicians, circulating nurses, and anesthesiologists or certified registered nurse anesthetists. Prior to study initiation, every participant signed an informed consent and was subsequently assigned a study identifier to maintain participant anonymity. In addition, participants filled out a demographic questionnaire that included information regarding their age, gender, role in the robotic OR and their years of experience in that role.

At the end of each surgery, all team members evaluated the quality of communication, performance, teamwork, and mental load using a 14-item questionnaire based on previously published validated questionnaire (11, 12). Agreement with the statement presented in each item was graded on a 10-point Likert scale (Table S1).

In the first phase of the study, the control period, team members did not use the headset device during surgery. Following accrual of adequate cases (see power calculations) for baseline data, the same participants proceeded to the second phase of the study, the intervention period, which involved the use of the headset device by all team members during the procedure. Through both phases of the study, participants filled out the questionnaire.

The Quail Digital Healthcare headset system that was used is a digital audio hands free, wireless headset device with noise cancellation capabilities that weighs 23 gm (Quail Digital, London, UK) (Figure 1).

**Figure 1.**
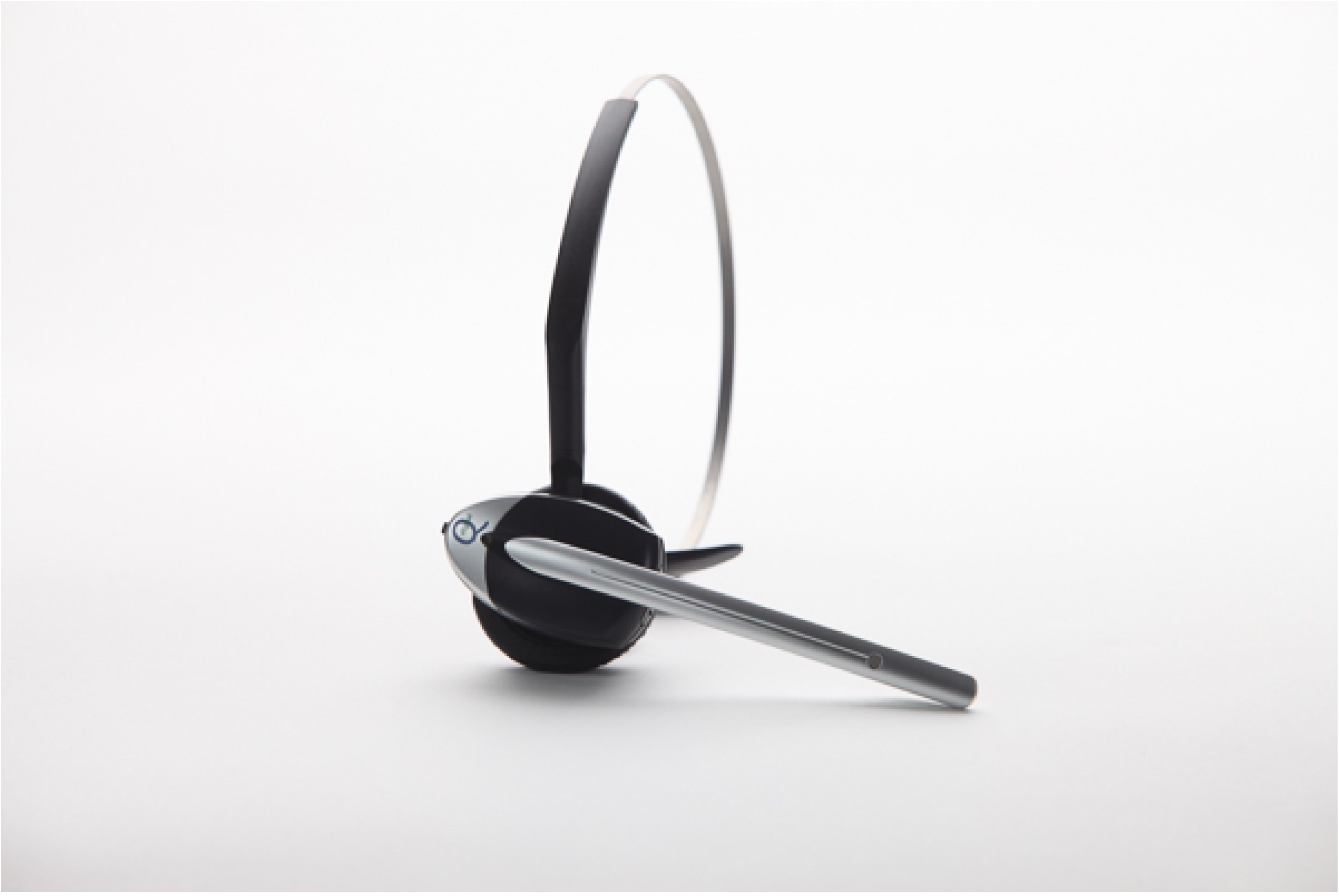
Quail digital wireless headset.

In addition, decibel-recording devices (TES-1352H, Taipei, Taiwan) were used during each case to monitor the noise level in the OR in both phases of the study. In each OR, 2 such devices were installed: 1 at the bedside (B), and 1 at the surgeon’s console (C). Ambient OR noise (dB) were continuously recorded and stored in 1-second intervals. Furthermore, background noise readings were taken once from each OR at the end of the trial. The background noise levels were subtracted from OR recordings.

The following patient clinical data were retrospectively retrieved from the electronic medical records: demographics, past medical and surgical history, body mass index, indication for surgery, type of surgery, time to complete the surgery, perioperative and postoperative complications, length of stay and pathology findings. The Charlson scoring system was used to summarize the comorbidity of patients in both the control and study phases (13). The severity of operative complications was graded according to the Clavien-Dindo classification (14). Patient outcome data were correlated with the survey and noise data obtained from study participants.

### Statistics

#### Questionnaire scoring

To summarize the questionnaire, an overall score was calculated. Scores from statements presented in a negative tone, e.g. “I had to repeat myself because people didn’t understand/hear my message the first time,” were subtracted from 10, so that all items could contribute in the same direction towards the overall score; i.e. higher overall scores indicated better communication and efficiency.

#### Processing of noise data

An average background noise level per room was calculated from each microphone in 6 different operating rooms. This average was subtracted from each noise time series from the surgical cases matched to the corresponding microphone and OR. The average noise and the percentage of time that noise surpassed 70 dB during console operation were calculated for each microphone in each case. These 2 measures were compared between the control and intervention groups using a linear regression model to account for the potential influence of the OR and team size.

#### Comparison of variables

Descriptive statistics were calculated and compared between groups. For patient characteristics the t-test and chi-square or Fisher exact tests were used to compare continuous and categorical variables respectively. The effect of headset use on patient outcomes was compared using a linear regression model that accounted for relevant patient characteristics. A linear mixed model was used for comparisons of individual item scores and overall score from the questionnaires between the groups to account for the clustered nature of the data. Significance was set at 0.05 for all comparisons.

#### Sample size calculation

Sample size for this trial was estimated considering operative time as the main outcome. A minimum of 50 procedures per arm were deemed necessary in order to detect a standardized difference of 0.50 in operative time between the 2 groups using a two-sided test at 80% power and significance level of 0.05. Based on historical schedules, such a size could be achieved by a 1-month period for each phase of the study.

## Results

A total of 137 robotic procedures performed from January through March 2015 were included in the analysis. The study group (with headsets) contained 69 cases and the control (without headsets) contained 68 cases.

Baseline characteristics and procedure details are presented in Table 1. No significant differences were found between the patients in the study and control groups in terms of body mass index, prior comorbidities, past surgical history, type of robotic surgery performed, use of the fourth robotic arm, and number of team members participating in the procedure.

**Table 1.** Descriptive statistics of patients and procedures.

| **Variable** | **With headsets**<br>**Mean ± SD**<br>**(n=69)** | **Without headsets**<br>**Mean ± SD**<br>**(n=68)** | **p-value** |
| --- | --- | --- | --- |
| Age (years) | 58.6 ± 11.0 | 58.1 ± 12.1 | .827 |
| Body mass index (lb/inch<sup>2</sup>) | 30.4 ± 5.7 | 30.7 ± 6.4 | .775 |
| **Variable** | **Counts (%)** | **Counts (%)** | **p-value** |
| Sex: Males | 42 (61%) | 41 (60%) | 1.000 |
| Department: Gynecology | 22 (32%) | 20 (29%) | .853 |
| Charlson score |  |  | .892 |
| 0 | 35 (51%) | 40 (59%) |  |
| 1 | 20 (29%) | 10 (15%) |  |
| 2 | 14 (20%) | 18 (26%) |  |
| Previous laparotomy |  |  | .674 |
| Single | 11 (17%) | 15 (22%) |  |
| Multiples | 4 (6%) | 5 (7%) |  |
| Previous laparoscopy | 12 (19%) | 13 (19%) | 1.000 |
| Procedure |  |  | .504 |
| Prostatectomy | 31 (45%) | 31 (46%) |  |
| Hysterectomy | 21 (30%) | 16 (24%) |  |
| Partial nephrectomy | 6 (9%) | 11 (16%) |  |
| Radical nephrectomy | 8 (12%) | 5 (7%) |  |
| Other<sup>a</sup> | 3 (4%) | 5 (7%) |  |
| LND performed | 25 (45%) | 26 (50%) | .701 |
| Use of a fourth robotic arm | 41 (59%) | 35 (51%) | .392 |
| Frozen sections | 31 (45%) | 27 (40%) | .605 |
| Surgical team size |  |  | .654 |
| 5 members | 5 (7%) | 4 (6%) |  |
| 6 members | 18 (26%) | 11 (16%) |  |
| 7 members | 14 (20%) | 17 (25%) |  |
| 8 members | 14 (20%) | 20 (29%) |  |
| 9 members | 14 (20%) | 12 (18%) |  |
| 10 members | 4 (6%) | 4 (6%) |  |

SD, standard deviation; LND, lymph node dissection
<sup>a</sup>Other: Study group - Gynecology: 1 myomectomy, Urology - 2 simple prostatectomies;
Control group - Gynecology: 1 sub-total hysterectomy and 2 myomectomies, Urology - 2
radical nephroureterectomy.

A total of 148 team members participated in the study. Demographic data and work experience of all participants, stratified by role, are summarized in Table 2. Overall 843 questionnaires were filled out, with a response rate of 89%. There was no significant difference in the response rate between the different team member categories (Figure 2).

**Table 2.** Participants demographics^a^.

| Characteristic | Attending | Anesthesia<sup>b</sup> | Circulator | ST | Fellow | Resident | FA |
| --- | --- | --- | --- | --- | --- | --- | --- |
| N | 13 | 54 | 21 | 11 | 5 | 20 | 4 |
| Age (years) | 49.7 ± 7.3 | 41.1 ± 11.4 | 45.8 ± 11.2 | 43.0 ± 12.9 | 38.0 ± 5.3 | 30.9 ± 2.4 | 40.0 ± 4.1 |
| Sex |  |  |  |  |  |  |  |
| Females | 2 (15.4) | 35 (64.8) | 19 (90.5) | 10 (90.9) | 1 (20.0) | 11 (55.0) | 4 (100.0) |
| Males | 11 (84.6) | 19 (35.2) | 2 (9.5) | 1 (9.1) | 4 (80.0) | 9 (45.0) | 0 (0.0) |
| Experience (years) | 16.9 ± 9.5 | 9.8 ± 10.3 | 15.6 ± 12.8 | 9.2 ± 12.8 | 5.0 ± 5.4 | 3.1 ± 1.5 | 10.0 ± 10.8 |
| Robotic experience |  |  |  |  |  |  |  |
| 0-20 cases | 1 (7.7) | 9 (16.7) | 5 (23.8) | 2 (18.2) | 0 (0.0) | 11 (55.0) | 0 (0.0) |
| 20-40 cases | 0 (0.0) | 5 (9.3) | 3 (14.3) | 2 (18.2) | 0 (0.0) | 5 (25.0) | 0 (0.0) |
| 40-60 cases | 2 (15.4) | 6 (11.1) | 2 (9.5) | 0 (0.0) | 0 (0.0) | 0 (0.0) | 0 (0.0) |
| 60-80 cases | 0 (0.0) | 7 (13.0) | 2 (9.5) | 2 (18.2) | 1 (20.0) | 0 (0.0) | 1 (25.0) |
| 80-100 cases | 0 (0.0) | 7 (13.0) | 1 (4.8) | 2 (18.2) | 0 (0.0) | 0 (0.0) | 0 (0.0) |
| > 100 cases | 10 (76.9) | 20 (37.0) | 8 (38.1) | 3 (27.3) | 4 (80.0) | 4 (20.0) | 3 (75.0) |

<sup>a</sup>128 of 148 participants provided the demographic information.174 <sup>b</sup>Anesthesia: Anesthesiologist/CRNA (certified registered nurse anesthetists)
Numbers represent mean ± standard deviation except for sex and robotic experience,
which represent counts (%). Age and sex are distributed significantly different at $p < .001$ ,177 experience $p = .001$ and robotic experience $p = .013$ .

**Figure 2.**
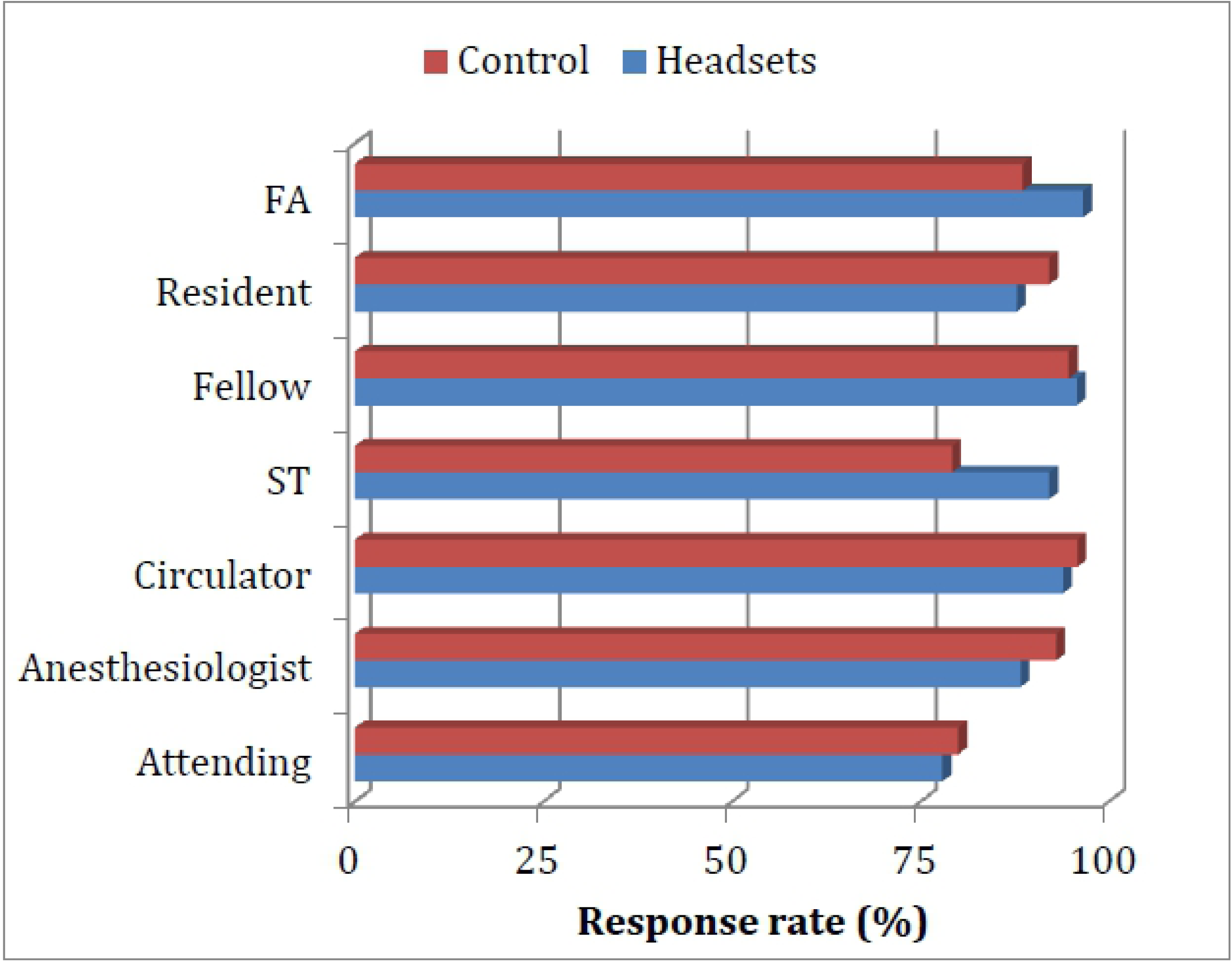
Response rate of team members, stratified by role. Abbreviations: FA, first assistant; ST, surgical technician.

Participants reported better overall communication in cases where headsets were used (113.0 ± 1.6 vs. 101.4 ± 1.6; p < .001) as measured by mean survey score. The mean score for individual items in both the study and control groups are shown in Table 3. When stratified by responder’s role, the overall scores were significantly higher while using the headsets, in comparison to the control. The mean score for each item with and without the headsets stratified by responder role is presented in Table S2.

**Table 3.** Mean scores for individual items of the survey^a^.

| **Variable** | **With headsets** | **Without headsets** | **p-value** |
| --- | --- | --- | --- |
|  | **Mean ± SE** | **Mean ± SE** |  |
| Heard clearly during case | 9.0 ± 0.2 | 7.2 ± 0.2 | <.001 |
| Needed to repeat | 7.6 ± 0.2 | 5.9 ± 0.2 | <.001 |
| Team communication | 8.6 ± 0.1 | 7.5 ± 0.1 | <.001 |
| Focus | 8.5 ± 0.1 | 8.0 ± 0.1 | <.001 |
| Steps took longer | 7.8 ± 0.2 | 6.3 ± 0.2 | <.001 |
| Feel safe | 8.9 ± 0.1 | 8.3 ± 0.1 | <.001 |
| Successful in task | 8.8 ± 0.1 | 8.5 ± 0.1 | <.001 |
| High team morale | 8.9 ± 0.1 | 8.4 ± 0.1 | <.001 |
| Participation | 8.9 ± 0.1 | 8.6 ± 0.1 | <.001 |
| Efficient teamwork | 9.0 ± 0.1 | 8.4 ± 0.1 | <.001 |
| Felt exhausted | 7.0 ± 0.2 | 6.3 ± 0.2 | <.001 |
| Felt stressed/irritated | 7.5 ± 0.2 | 7.0 ± 0.2 | <.001 |
| Hard work for task | 5.8 ± 0.2 | 5.3 ± 0.2 | <.001 |
| Noise bothered/distracted | 7.5 ± 0.2 | 6.8 ± 0.2 | <.001 |

SE, standard error
<sup>a</sup>For the full version of each statement please refer to Table S1.

Use of the headset did not reduce the noise level in the robotic OR, as there was no significant difference in average noise levels between the groups (Table 4). However, cases in which headsets were used demonstrated a lower percentage of time with noise level above 70 dB at the console (8.2% ± 0.6 vs. 5.3% ± 0.6, p < .001) (Table 4). There were no differences between the study and control groups in regards to time to complete the surgery, estimated blood loss, length of hospital stay, or postoperative complications (Table 4). Similarly, there were no differences in outcomes after controlling for procedures type.

**Table 4.**
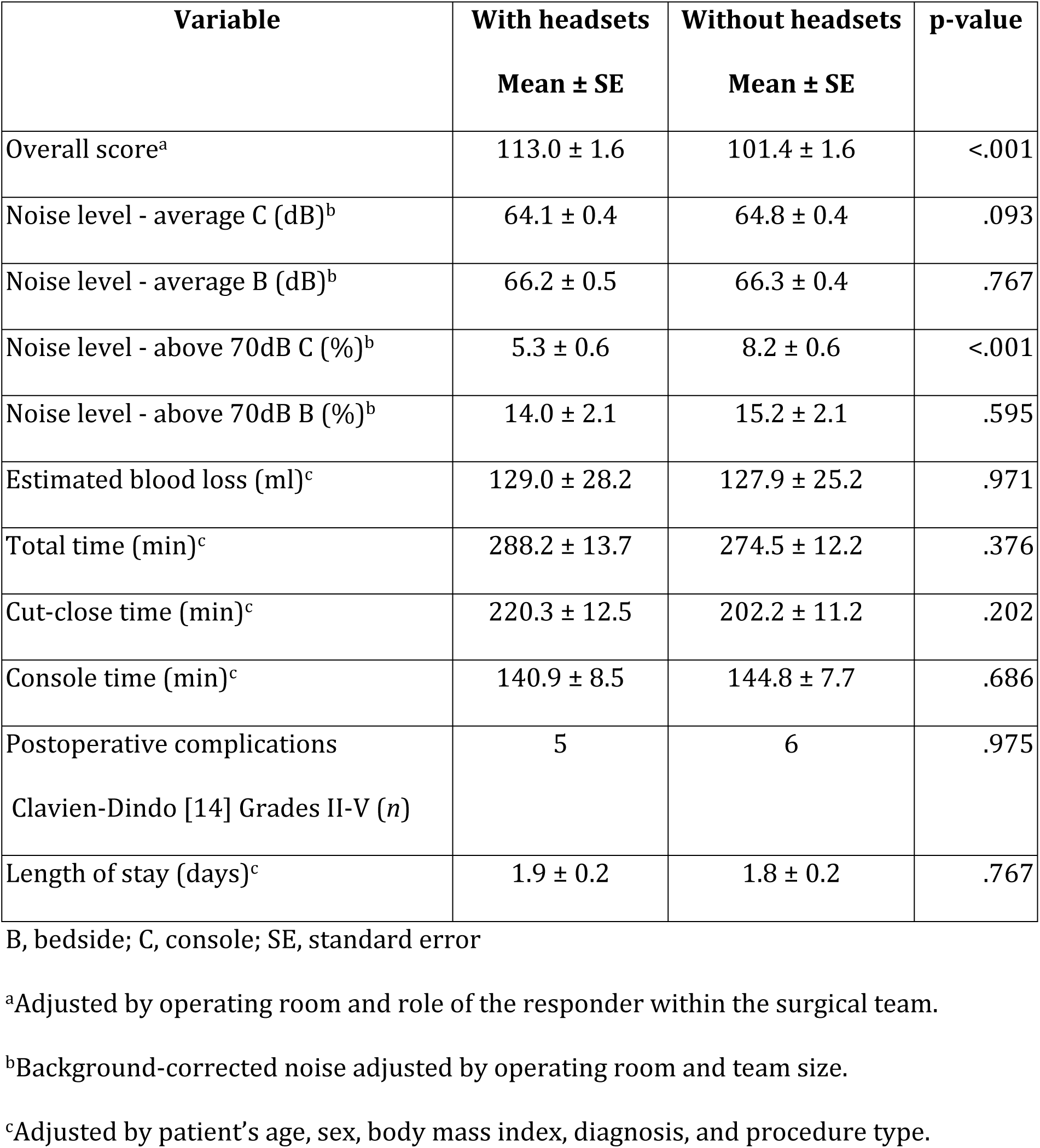
Outcomes comparison by use of headsets.

## Discussion

This study is the first to evaluate the impact of using wireless headsets on the quality of communication between different team members during robotic surgery. Our data shows that the use of such a device significantly improves the quality of communication in robotic surgery. However, these improvements did not reduce operation duration nor clinical outcomes for patients.

The last decade is characterized by a rapid adoption and dissemination of robotic technology and robotic-assisted minimally invasive surgery. In urology, for example, robotic-assisted radical prostatectomy was rapidly adopted to become the main surgical approach for treatment of localized prostate cancer, with an increasing rate from 8% in 2004 to 67% in 2010 (15). Between 2005 and 2010, the rate of robotic hysterectomy increased from as low as 0.5% to as high as 22% (16). It is estimated that approximately one-half of all minimally invasive hysterectomies are now performed robotically (17, 18). The rapid assimilation of the robotic platform is attributed to its hypothetical benefits, such as improved ergonomics, wider range of motion, 3-dimenssional stereoscopic vision and enhanced visual magnification. However, adoption of such technology raised concerns about patient safety and surgical outcomes (19-21).

The causal relationship between communication failure in the OR and adverse patient outcomes is well established in the literature (1-3). The Joint Commission identified breakdowns in communication as a leading root cause of operative or postoperative complication events (3). In their observational study, Wiegmann and colleagues found that surgical errors increased significantly with increased disruptions and that teamwork and communication problems were the strongest predictors of surgical errors (22).

Only four studies have looked at the inherent communication challenges faced by team members in robotic surgery. Robotic surgery was associated with a higher volume of information exchanged within the OR team, compared with regular laparoscopy during cholecystectomy (8). Specifically, there was a significant shift of communication in robotic OR towards verbal cues (6, 7).

Our prospective controlled trial is the first intervention aiming to improve communication and teamwork in the robotic OR. We focused on the most important part of communication in the robotic setting, which is verbal communication. With the assistance of the wireless headsets, we amplified the verbal cues. This intervention showed marked improvement in all 4 of the domains that were evaluated in the questionnaire: quality of communication, performance, teamwork, and mental load. Moreover, all team members, regardless of their role, shared the same positive perception of the added value of using the headsets in robotic cases.

Previous studies have found a correlation between the level of noise in the OR and postoperative complications (9, 10). Kurman et al. measured the sound level during 35 elective open abdominal procedures and found that increased intraoperative noise volume was associated with surgical site infection (9). In their prospective controlled trial, Engelmann and colleagues assessed the impact of a noise reduction program in a pediatric OR. Sound levels were measured and surgical complications recorded before and after implementation of a noise reduction protocol. The intervention significantly reduced the noise level in the OR. Additionally, the incidence of postoperative complications was significantly lower among the patients in the intervention group (10).

Our data showed that in cases where headsets were used there was a decreased period of time with noise level above 70 dB at the robotic console. This level of noise is equivalent to a domestic vacuum cleaner (23). However, there was no significant difference in the average noise level between both groups. Together, these data suggest that reduction in peak noise duration is not sufficient to improve patient outcomes and a reduction in the average noise levels is necessary.

Although noise was significantly reduced only at peak levels, our data demonstrated that team members perceived communication to be better while using the headsets.

Specifically, participants commented that they could hear clearly during the case, needed to repeat themselves less and were less bothered or distracted by the noise in the OR when using the device. This can be attributed to the noise cancelling capabilities of the headsets, which simultaneously reduces ambient noise and increases voice clarity for the user.

Our study has its own limitations. Firstly, this study was not randomized because the team members themselves were the study participants, and randomization was not feasible due to a lack of consistency of participants assigned to cases during the study period.

Additionally, the decision to start with the control phase first was based on the concern that team members could potentially be biased by the prior use of the headset device. Despite the use of validated questionnaires, our results may have been subject to responder bias. Participants who had a good experience with the headsets might be more enthusiastic to fill out the survey at the end of the case, as compared to participants who did not have a good experience with the headsets. This could potentially skew results in favor of this device. However, the fact that we had a response rate of 89% with no significant difference in the response rates by participant’s role suggests that such potential bias was negligible.

## Conclusion

We present here a novel approach to address a communication challenge in robotic procedures. Our study shows that the use of wireless headsets improves communication in the robotic OR. In addition, the percentage of time above a peak sound level of 70 dB is reduced while using headsets. These changes did not affect the clinical outcomes.

## Acknowledgments

We thank Dr. Amir Shafat, National University of Ireland Galway, for scientific discussion and editorial assistance. We thank Ms. Stephanie Stebens for editorial assistance.

